# Characterization and safety profile of Imuno TF^®^, a nutritional supplement for immune system regulation containing transfer factors

**DOI:** 10.1101/2020.12.07.414169

**Authors:** Hudson Polonini, Any Elisa de Souza Schmidt Gonçalves, Eli Dijkers, Anderson de Oliveira Ferreira

## Abstract

Imuno TF^®^ is a nutritional supplement composed of isolated transfer factors (TF) from porcine spleen. It is composed of a specific mixture of molecules that impact functions of the biological systems, and historically is linked to the immune system regulation. In this study, we demonstrate for the first time its proteomic analysis, nutritional composition, and safety profile in terms of mutagenic potential and acute oral dose (LD_50_). The obtained analysis indicated the product is a complex set of oligo- and polypeptides constituted of 163 different peptides which can potentially act on multiple mechanisms on the immune system pathways. The chemical composition showed low fat and low sugar content, saturated fatty acids-free and the presence of 10 vitamins and 11 minerals. No mutagenic effect was observed, and the LD_50_ was 5,000 mg kg^-1^ body weight. This accounts for a safe product to be used by oral route, with potential benefits for the immune system.

## 1. INTRODUCTION

Imuno TF^®^ is a nutritional supplement sold worldwide and composed of oligo- and polypeptides fractions from porcine spleen, commonly referred to as transfer factors (TF) and with biological activity on immune regulation. They are produced by T helper cells(Fudenberg & Fudenberg, 1989; Welch et al., 1976) and are non-species-specific, i.e., TF produced in one species is effective in another animal species.(Fudenberg & Fudenberg, 1989; Wilson et al., 1987)Although there is a number of evidences on its clinical effects,(Steele et al., 1980; Viza, Fudenberg, Palareti, Ablashi, Vinci, et al., 2013) to date few studies have evaluated its molecular characterization.

The first mention of TF in the literature was in 1955, where it was demonstrated that a dialysis of leukocyte extract from a healthy donor - which presents a positive immune response confirmed through delayed hypersensitivity tests - was able to transfer to a healthy recipient and to also respond positively to this test.(Lawrence, 1955) In 1983, Lawrence and Borkowsky modified the original TF purification protocol using a dialysis membrane and a second molecular exclusion membrane to obtain molecules of three different sizes: <3.5 kDa, >3.5 kDa, and <12 kDa. This allowed them to identify the fraction that had the ability to bind to antigens: 3.5 kDa and 12 kDa.(Lawrence & Borkowsky, 1983) In subsequent studies, Kirkpatrick and collaborators characterized these molecules as being small peptides, with molecular weight generally between 3.5 kDa and 6.0 kDa.(Kirkpatrick, 1993; Rozzo & Kirkpatrick, 1992)

Since then, more studies have attempted to shine light on the molecular structure of TF. There is evidence that TFs consist of short chains of amino acids with small pieces of ribonucleic acid (RNA) attached.(Berrón-Pérez et al., 2007; Kirkpatrick, 1996) The RNA is probably related to a cytophilic property and the specificity of the TF(Krishnaveni, 2013), as the absence of the oligoribonucleotide linked to the amino termination of peptide results in loss of activity.(Viza, Fudenberg, Palareti, Ablashi, De Vinci, et al., 2013)

The length of the peptide chain of TFs, is still under debate; there are reports of 17 or 18 amino acids present in TF,(Guilan et al., 1990; Liu et al., 2007) with high tyrosine and glycine content,(Krishnaveni, 2013) but given the molecular weight of the fractions, they could be even bigger molecules with 24 amino acids or more. Especially since, the molecular weight of tryptophan (the heaviest amino acid) for example, is more than 200 Da.(White, 2009)

It also needs to be taken into consideration that by nature, TFs extracts are complex mixtures, containing a high number of different TFs, and not just one single chemical entity.(Hennen, 1998) In contrary to Imuno TF^®^, other commercially available traditional TF extracts are generally obtained from cow colostrum (which can cause allergic reactions in other species due to the presence of immunoglobulins(Bernhisel-Broadbent et al., 1991)), bird’s egg yolks, or other tissue obtained from suitable animals.(Hennen, 1998; Krishnaveni, 2013) Until now, little is known about the exact chemical composition of the TFs.

The objetive of this study was to characterize Imuno TF^®^ though proteomics analysis and to evaluate its safety profile through the determination of the mutagenic potential and acute oral toxic dose. Imuno TF^®^ was chosen as it is derived from porcine spleen and therefore has a more favorable safety and efficacy profile, compared to TFs from other sources, which may contain immunoglobulins.(Bernhisel-Broadbent et al., 1991)

## 2. MATERIAL AND METHODS

### 2.1. Samples

To ensure meaningful results on chemical composition, samples (30 g) from three different batches of powdered Imuno TF^®^ were supplied by Infinity Pharma Brasil (Campinas, SP, Brazil), a Fagron company (Rotterdam, The Netherlands).

### 2.2. Nutritional composition (Compositional analysis)

The nutritional composition of Imuno TF^®^ was assessed through different methods: monounsaturated, polyunsaturated and *trans* fats were determined by gas chromatography (GC), and total dietary fiber by gravimetry; cholesterol, carbohydrates (fructose, glucose, maltose, lactose, and sorbitol), amino acids and vitamins were determined by high performance liquid chromatography (HPLC) and protein by titrimetry; vitamin B12 was determined by enzyme-linked immunosorbent assay (ELISA); starch and sucrose by UV spectrophotometry; and mineral composition content was determined by Inductively Coupled Plasma mass spectrometry (ICP-MS).

### 2.3 Proteomic analysis

#### Protein extraction

Imuno TF^®^ samples (n=3) were solubilized in ammonium bicarbonate and digested in a 1:80 trypsin solution for 8h. Resulting peptides were analyzed using a two-dimensional Acquity M-Class nanoUPLC system (Waters Corporation, Milford, MA), coupled to a Synapt G2-Si spectrometer (Waters Corporation, Milford, MA). The MS and MS/MS data were obtained using data-independent acquisition (DIA) and ion mobility separation. The peptide samples derived from the three biological and technical triplicates (in 1% formic acid) were analyzed using an HDMSE (high-definition data-independent mass spectrometry) employing a 54 min gradient in reverse-phase chromatography (linear gradient of 3–40% ACN).

#### Protein identification and MS analysis

Peptides were loaded onto a nanoACQUITY UPLC HSS T3 Column (100 A, 1.8 μm, 75 μm × 150 mm, Waters Corporation, Milford, MA, United States). Injections were performed using a nano-electrospray ionization source in positive ion mode (nanoESI (+), with a NanoLock-Spray (Waters, Manchester, United Kingdom) ionization source. A solution of [Glu1]-Fibrinopeptide B (Glu-Fib; Human) was used as the lock mass and was sampled every 30 seconds.

MS and MS/MS spectra were processed, and data searched in Progenesis QI for Proteomics 3.0 (Waters). In this process, the following parameters were considered: 5 as maximum peptide charge, maximum protein mass of 600 kDa, a maximum of 2 missed cleavages, at least 2 fragments per peptide, with a False Discovery Rate (FDR) FDR of < 1% and a mass error cutoff of 20 ppm. The tryptic digestion of a low molecular mass fraction can generate peptides with only one tryptic end; thus, data was searched using trypsin, Lys-C and Arg-C as enzymes. Cysteine carbamidomethylation and methionine oxidation were considered as fixed and variable modifications, respectively. The database used was the *Sus scrofa* (Pig) UNIPROT databank.

### 2.4 In vitro safety profile: mutagenic potential

#### Sample and controls

Imuno TF^®^ was diluted in culture medium (CM) [DMEM, containing penicillin, streptomycin, L-glutamine, sodium pyruvate and 5% fetal bovine serum (FBS)] to 100; 10; 1; 0.1; 0.01; 0.001; 0.0001 and 0.00001 mg mL^-1^ for cell viability and at 5, 1.67 and 0.56 mg mL^-1^ (with and without S9 metabolization) for micronucleous testing.

Sodium dodecyl sulfate (SDS) was used as positive control (starting concentration = 100 µg mL^-1^ in CM; seven solutions were prepared from the initial concentration, in dilution factor 1.47) for cell viability and cyclophosphamide (20 µg mL^-1^ in CM + 5% FBS; in the tests with and without S9 metabolization) and colchicine (0.1 µg mL^-1^, in CM + 5% FBS; long-term treatment, without metabolization) were used for the micronucleus test. CM was used as negative control for both tests.

#### Cell viability

Cell viability test was conducted following the OECD guidelines(OECD - Organisation for Economic Co-operation and Development, 2016) in order to identify the non-cytotoxic concentrations of Imuno TF^®^ to be used in the micronucleus test. The evaluation was – performed using the method of sulforhodamine B and the protein quantification was determined spectrophotometrically. Briefly, lung cells of the Chinese hamster (V79-4) were incubated in DMEM culture medium containing 10% of FBS and were kept under CO_2_ atmosphere for approximately 24 hours. After the adaptation period, the cells were exposed to 8 concentrations of Imuno TF^®^ for a period of 24 hours. After this period, cells were submitted to morphologic analysis; then, the cells were fixated with 50% (v/v) trichloroacetic acid followed by refrigeration for 1 hour. The plates were submitted to five washings in running water for the removal of the trichloroacetic acid residues, culture medium, FBS and secondary metabolites. After complete drying, the protein sulforhodamine B colorant was added at 0.4% (p/v) dissolved in acetic acid at 1% (v/v) and then, the plates were incubated at room temperature, for 10 minutes. The wells were washed 5 times with acetic acid solution 1% (v/v) and, after the complete drying, the colorant connected to the cell proteins was solubilized with Tris Base solution 10 mM. Spectrophotometric reading at 515 nm were obtained for the determination of the IC_50_ using XLSTAT 2020 software.

#### In vitro micronucleus test (MNvit)

V79-4 cells were incubated in culture medium (DMEM, L-glutamine, sodium pyruvate and 10% FBS) for around 24 hours. After this period, the cells were treated with three concentrations of Imuno TF^®^ for 3 hours, split into 2 groups: with or without S9-mix solution (MgCl_2_/KCl solution; glucose 6-phosphate; NADP; S9 fraction and reverse osmosis water). After the treatment period, the culture medium was removed, and the cells from the group without the S9-mix solution were exposed to the cytochalasin B solution (3 µg mL^-1^ in culture medium) for 21 h in the CO_2_ incubator. After this period, cells suspensions were obtained by centrifugation and the pellet was resuspended in refrigerated hypotonic solution for 3 minutes. Then, methanol:acetic acid (3:1) fixative was added; the suspensions were centrifuged and the pellet was resuspended in the fixative and again centrifuged – this step was then repeated one more time. The supernatant was discarded, leaving around 1 mL of fixative to resuspend the pellet. The cell suspension was dripped in two histological slides, previously cleaned, and kept in refrigerated reverse osmosis water. After drying, slides were cut with blue methylene eosin colorant (Giemsa/methanol) for five minutes. After a quick washing in reverse osmosis water and drying of the slides, the optical microscopy analysis was started and of each Imuno TF^®^ concentration tested, 2,000 binucleated cells were counted. In this population, the incidence of multinucleated cells with 1, 2, 3 or 4 micronuclei was considered. In addition, the number of mononucleated and multinucleated cells was accounted. With these experimental values, the CBPI (Cytokinesis-Block Proliferation Index), and the RI (Replication Index) were calculated, both to measure the cytotoxicity of the test items, and the percentage of binucleated cells with micronucleus (%BCMN) for the evaluation of the mutagenic potential. Data obtained from the slides was analyzed using XLSTAT software, and an ANOVA followed by a post-test Tukey was conducted to analyze the difference between the treatments (p < 0.05).

### 2.5. *In vivo* safety profile: median lethal dose (LD_50_)

#### Animals

Six healthy young female (nulliparous and not pregnant) Wistar Hannover line (Rattus Norvegicus) rats, with a weight range of approximately 200 g were used. Upon arrival at the laboratory, animals were evaluated, randomized, placed in cages with 5 animals per sex and allowed to acclimate for 5 days. During this period, animals were observed by a veterinary for general health and mortality. During the 24 hours duration of the experiment, animals were housed individually. This experiment was conducted following the OECD guidelines(OECD - Organisation for Economic Co-operation and Development, 2001) and supported by Protocol No. 004/20, approved by the Ethics Committee on the Use of Animals (CEUA) on 06/04/2020.

#### Acute oral dose

Animals were tested in a stepwise procedure with three female animals per step. A starting dose of 2,000 mg kg^-1^ of body weight (bw) was used. Before administration of Imuno TF^®^, animals were fasted over-night by depriving food; water was supplied normally. After 3-4 hours of Imuno TF^®^ administration, animals were allowed food again.

Imuno TF^®^ was diluted in water and administered as single dose by oral gavage as a single dose. The treated animals revealed no clinical signs on Day 1 and no mortality was observed. Then, a confirmation study (with again single dose of 2,000 mg kg^-1^ bw Imuno TF^®^) was ran in another 3 rats, 24 h after the first dose had been administered. After dosing, all animals were evaluated at least once during the first 30 minutes, and then periodically during the first 24 hours, with special attention during the first 4 hours after administration. After 24 h, the animals were evaluated daily for a period of 14 days. All changes observed during this period were systematically recorded, and individual records were maintained for each animal. The observations included: changes in the skin and hair, eyes, and mucous membranes and in the respiratory, circulatory, autonomic and central nervous system, somatomotor activity and behavioral pattern. Special attention was given to the appearance of tremors, convulsion, salivation, diarrhea, lethargy, drowsiness or coma. If any sign of suffering and stress was observed, animals were euthanized. In cases of euthanized animals, as well as in the case of dead animals, the time of death was precisely recorded. The animals were weighed before the administration of the test item (Day 0) and at days 7 and 14. At the beginning of the experiment, the body weight variation must not exceed 20% of the weight average.

At the end of the study, the surviving animals were euthanized in a CO_2_ chamber. A second method of euthanasia (cervical displacement) was performed to confirm the death of the animal. All animals were submitted to necropsy whenever possible. This included a careful examination of the outer surface of the body, all orifices, chest, pelvic and abdominal cavity, and their contents. The observed macroscopic changes were recorded. For a more detailed verification of the toxicity, autopsy was performed, and their organs were extracted for histopathological evaluation. The collected organs were stored in 10% formalin and then slides for evaluation were made. The organs collected were brain, spinal cord, stomach, large and small intestine, liver, kidneys, adrenals, spleen, lung, heart, thymus, ovaries, vagina, bladder and lymph node). Any observed microscopic changes were recorded.

## 3. RESULTS

### 3.1. Nutritional composition

The nutritional composition of Imuno TF^®^ can be found in **Table 1**.

**Table 1.** Nutritional composition of Imuno TF^®^.

| *Energy* | *Value per 100 mg* | *Reference intake*(European Commission, 2011) |
| --- | --- | --- |
| Energy value | 0.321 Kcal / 1,347 KJ | 2,000 Kcal |
| *General Components* | *Amount (mg per 100 mg)* | *Reference intake* |
| Total fat | < 0.3 | 70 g |
| of which |  |  |
| – saturated fat | < 0.01 | 20 g |
| – monounsaturated fat | < 0.01 | - |
| – polyunsaturated fat | < 0.01 | - |
| – *trans* fat | < 0.002 | - |
| – cholesterol | < 0.002 | - |
| Total carbohydrates |  | 260 g |
| of which |  |  |
| – starch | < 0.09 | - |
| – fructose | < 0.1 | - |
| – glucose | < 0.1 | - |
| – sucrose | < 0.1 | - |
| – maltose | < 0.1 | - |
| – lactose | < 0.2 | - |
| – sorbitol | < 0.1 | - |
| Fiber | 0.5 | - |
| Protein | 63.97 | 50 g |
| of which (amino acids) |  |  |
| – aspartic acid | 2.95 | - |
| – glutamic acid | 5.19 | - |
| – serine | 1.89 | - |
| – histidine | 0.93 | - |
| – glycine | 3.06 | - |
| – threonine | 1.41 | - |
| – arginine | 4.16 | - |
| – alanine | 0.50 | - |
| – tyrosine | 1.83 | - |
| – cystine | < 0.1 | - |
| – valine | 1.30 | - |
| – methionine | 1.17 | - |
| – phenylalanine | 0.06 | - |
| – isoleucine | 0.96 | - |
| – leucine | 1.39 | - |
| – lysine | 3.49 | - |
| – proline | 0.10 | - |
| Salt (sodium) | 0.7 | 6 g |
| ***Specific Components*** | ***Amount (per 100 mg)*** | ***Nutrient Reference Value (NRV)*** |
| Vitamins and minerals |  |  |
| Vitamin A | < 0.0005 µg | 800 µg |
| Vitamin B1 | < 0.0004 mg | 1.1 mg |
| Vitamin B2 | 0.00074 mg | 1.4 mg |
| Vitamin B3 | 0.00002 mg | 16 mg |
| Vitamin B6 | 0.00645 mg | 1.4 mg |
| Vitamin B9 | < 0.42 µg | 200 µg |
| Vitamin B12 | < 0.004 µg | 2.5 µg |
| Vitamin C | 0.02 mg | 80 mg |
| Vitamin E (alfa-tocopherol) | < 0.0002 µg | 12 mg |
| Vitamin E (gamma-tocopherol) | < 0.00035 mg | - |
| Potassium | 1.91 mg | 2,000 mg |
| Calcium | 0.028 mg | 800 mg |
| Phosphorous | 0.72 mg | 700 mg |
| Magnesium | 0.078 mg | 375 mg |
| Iron | 0.099 mg | 14 mg |
| Zinc | 0.0093 mg | 10 mg |
| Copper | 0.0065 mg | 1 mg |
| Manganese | 0.00011 mg | 2 mg |
| Selenium | 0.0011 mcg | 55 mcg |
| Chromium | < 0.000005 mcg | 40 mcg |
| Molybdenum | 0.019 mcg | 50 mcg |

Reference intake of an average adult (8,400 kJ/2,000 kcal).

### 3.2. Proteomic analysis

The characterized proteins identified on Imuno TF^®^ are described in **Table 2**. The proteomic analysis revealed a total of 163 peptides associated to 23 distinct *Sus scrofa* (Pig) proteins. Two out of the 23 identified proteins are still uncharacterized, meaning the existence of these proteins were predicted *in silico* and no functional studies have been conducted thus far. Peptides composition varied from 4 to 34 amino acids. The protein (or peptides from protein) with the highest prevalence in the sample are hemoglobin and globin domain (**Table 2**). Additionally, peptides from proteins which are known to interact with the immune system were identified: Talin-1, Ubiquitin-40S ribosomal protein S27a (aka 40S ribosomal protein S27a) and Ubiquitin-60S ribosomal protein L40.

**Table 2.** Proteomic analysis of Imuno TF^®^.

| Proteome identifier (UPID) | Gene name | Description |
| --- | --- | --- |
| P01965 | HBA | Hemoglobin subunit alpha |
| P02067 | HBB | Hemoglobin subunit beta |
| P02067 | HBB | Hemoglobin subunit beta |
| F1SFZ8 | TLN1 | Talin-1 |
| A0A287AZA7 | RPS27A | 40S ribosomal protein S27a (Ubiquitin-40S ribosomal protein S27a) |
| A0A5K1UHC4 | AGPAT5 | PlsC domain-containing protein |
| P01965 | HBA | Hemoglobin subunit alpha |
| Q06AT1 | HPCA | Neuron-specific calcium-binding protein hippocalcin |
| A0A480JNE4 | FAM184A | Protein FAM184A isoform 1 |
| P49756 | RBM25 | RNA-binding protein 25 |
| P09571 | TF | Serotransferrin |
| Q8WXA9 | SREK1 | Splicing regulatory glutamine/lysine-rich protein 1 |
| Q96A84 | EMD1 | EMI domain-containing protein |
| A0A287AAR0 | GIMAP7 | AIG1-type G domain-containing protein |
| A0A287AHD6 | LOC100622504 | Uncharacterized protein |
| P00819 | ACYP2 | Acylphosphatase-2 |
| P63053 | UBA52 | Ubiquitin-60S ribosomal protein L40 |
| P13796 | LCP1 | Plastin-2 |
| A0A5G2Q920 | ENSSSCG00000049439 | Uncharacterized protein |
| Q99873 | PRMT1 | Protein arginine N-methyltransferase 1 |
| A0A287AL08 | TAFA2 | Protein FAM19A2 isoform X1/ Chemokine-like protein TAFA-2 |
| A0A5G2QDH3 | ENSSSCG00000051012 | Reverse transcriptase domain-containing protein |
| F1SVA9 | SH2D6 | SH2D6 (signal transductor) |

### 3.3. In vitro safety profile: mutagenic potential

Cell viability was determined in terms of half maximal inhibitory concentration (IC_50_). The IC50 value found for Imuno TF^®^ was 11.6 mg mL^-1^. From the results of the cell viability test, 0.56, 1.67 and 5.0 mg mL^-1^ were selected for the evaluation of the mutagenic potential in the in vitro micronucleus test. In The pH obtained for the highest concentration was 7.5. Positive control (SDS) presented IC_50_ was equal to 39.6 µg mL^-1^ with a pH of 7.5.

In addition to the viability test, the cytotoxic and cytostatic activity of the treatments in relation to the negative control items was evaluated from the values of CBPI and RI (Table 3). The CBPI indicates the mean number of cycles that each cell suffers during an exposure period to cytochalasin B and may be used to calculate the cell proliferation. In addition, the RI indicates the relative number of nuclei in treated cells, in comparison with the negative control item of the cultures and may be used to calculate the percentage of cytostatic cells that correspond to the inhibition of cell growth. Thus, the RI is a form of comparison of the number of binucleated or multinuclear cells that are found in the division process and the higher its value, the smaller is the number of cytostatic cells, consequently, the lower will be the cytotoxicity of the test item.

**Table 3.** Values concerning the CBPI, RI and %BCMN of Imuno TF^®^ after the 3-hour and 24-hour treatments.

| Test items |  | 3h, S9- |  |  | 3h, S9+ |  |  | 24h, S9- |  |  |
| --- | --- | --- | --- | --- | --- | --- | --- | --- | --- | --- |
| Name | Concentration | CBPI | RI (%) | %BCMNI | CBPI | RI (%) | %BCMNI | CBPI | RI (%) | %BCMNI |
| Negative control | 100% | 1.93 ± 0.002 | 100.00 ± 0.00 | 1.85 ± 0.07 | 1.91 ± 0.016 | 100.00 ± 0.00 | 2.00 ± 0.00 | 1.94 ± 0.020 | 100.00 ± 0.00 | 1.85 ± 0.07 |
| Positive control | 20 µg mL<sup>-1</sup> | 1.92 ± 0.004 | 99.25 ± 0.74 | 1.85 ± 0.07 | 1.90 ± 0.002 | 98.84 ± 1.92 | 5.00 ± 0.14* | 1.66 ± 0.005 | 70.87 ± 0.97 | 6.55 ± 0.35* |
| Imuno TF<sup>®</sup> | 5 mg mL<sup>-1</sup> | 1.93 ± 0.008 | 99.65 ± 0.65 | 2.15 ± 0.07 | 1.89 ± 0.004 | 98.20 ± 2.16 | 2.15 ± 0.07 | 1.85 ± 0.022 | 90.93 ± 0.35 | 2.45 ± 0.07 |
|  | 1.67 mg mL<sup>-1</sup> | 1.93 ± 0.002 | 99.94 ± 0.05 | 2.05 ± 0.07 | 1.92 ± 0.007 | 101.11 ± 0.98 | 2.05 ± 0.07 | 1.91 ± 0.007 | 97.26 ± 1.35 | 2.25 ± 0.07 |
|  | 0.56 mg mL<sup>-1</sup> | 1.93 ± 0.003 | 99.94 ± 0.57 | 1.85 ± 0.07 | 1.91 ± 0.006 | 99.81 ± 1.16 | 1.95 ± 0.07 | 1.93 ± 0.007 | 98.75 ± 1.39 | 2.05 ± 0.07 |

\*p<0.05 in relation to negative control item. Results expressed as mean of the replicates ± standard deviation; Negative Control Item: cells containing only culture medium; Positive Control Items: Cyclophosphamide (3h) and Colchicine (24h); S9-: without S9 metabolization; S9+: with S9 metabolization; CBPI: Cytokinesis-Block Proliferation Index; RI: Replication Index; %BCMNI: percentage of binucleated cells with micronucleus.

The CPBI and RI values of the concentrations of the test item and the positive control item were within the acceptance criteria defined by the test guide for both the 3-hour treatment and the 24-hour treatment. In the mutagenicity test, CBPI and RI values were above 1.6 and 70%, respectively, confirming the absence of cytotoxicity in the concentrations of the evaluated items.

For the experimental model with metabolic activation, the post-mitochondrial fraction (S9 fraction) was included. The results of BCMN percentage obtained with the use of the S9 fraction (S9+) combined with those of the experiment without S9 (S9-) allow to determine if the Imuno TF^®^ tested is mutagenic in its original form (percentage values of BCMN similar in both experiments), if it becomes mutagenic after metabolization (values of BCMN S9+ are higher than BCMN S9-) or if generated metabolites are less mutagenic (values of BCMN S9-are higher than BCMN S9+).(OECD - Organisation for Economic Co-operation and Development, 2016) For Imuno TF^®^, it was shown that both in systems with and without S9 metabolization, there was no statistically significant increase in the percentage of MN (Table 3) for the duration of 3 or 24 hours.

### 3.4 In vivo safety profile: median lethal dose (LD_50_)

Table 4 shows the weights of animals (g) during the study and the mortality index. No weight loss was observed in the tested animals during the study period. No macroscopic finding was observed after necropsy, both on external and visceral examination of terminally sacrificed rats. The microscopic evaluation revealed preserved organs and no abnormalities. It must be considered that the analyzed tissues did not show signs of toxic lesions, necrotic conditions or inflammatory lesions (tissues preserved before the morphological evaluation). In the absence of any pathological injury in terminally sacrificed rats, it is concluded that the test item did not produce any treatment related effect at the dose level.

**Table 4.** Weights of animals and mortality index for the Imuno TF^®^ evaluation of the oral acute-dose toxicity.

| Experiment | Animal number | Animal weight (g) |  |  | Mortality index (%) |
| --- | --- | --- | --- | --- | --- |
|  |  | Day 0 | Day 7 | Day 14 |  |
| 1 | 1 | 225 | 240 | 251 | 0.0* |
|  | 2 | 204 | 230 | 237 |  |
|  | 3 | 185 | 197 | 207 |  |
| 2 | 1 | 167 | 185 | 192 | 0.0* |
|  | 2 | 191 | 211 | 229 |  |
|  | 3 | 202 | 222 | 243 |  |

\*No clinical sign was observed in any animal during the 14-days period, nor pathologic (macroscopic) or histopathologic (microscopic: toxic lesions, necrotic conditions or inflammatory lesions) findings on skin, brain, eyes, lungs, heart, liver, spleen, urinary system, intestines, reproductive tract and body as a whole. Dose of 2,000 mg kg<sup>-1</sup> bw.

## 4. DISCUSSION

Stimulation of the immune response, without disturbing it’s function can be obtained by the use of immunologically active substances localized in the lymphoid tissues and/or non-lymphoid origin, particularly those in the spleen. The spleen is the source of a large amount of biologically active substances, as well as protein factors which can enhance the humoral immune response and it’s inhibition.(Zaiko et al., 2014)

Imuno TF^®^ is an ultrapurified extract from porcine spleen, and its nutritional composition shows that it contains at least 10 different types of vitamins and 11 minerals, from which two have well-stablished roles contributing to the normal function of the immune system (selenium and zinc).(EFSA - European Food Safety, 2009; European Food Safety Authority, 2011) It is also a supplement with high content of protein and low quantities of fat and carbohydrates, and therefore can be considered a low fat, saturated fatty acids-free and low sugar product.(European Commission. Directorate General Health and Consumer Protection, 2001)

Understanding the composition of this nutrient allows us to associate the characterization with its biological function. The amino acids found in Imuno TF^®^ are composed from at least 163 peptides, indicating that Imuno TF^®^ is a complex set of peptides. Possibly, this is even an underestimation. A higher number could be present, as the high abundance of some peptides can mask the presence of other’s occurring in smaller quantities. As expected, a product derived from spleen presents also a considerable amount of cell debris or blood-derived proteins. Each peptide identified appears to be composition ranging from 4 to 34 amino acids (after digestion for the analysis), but bigger chains are expected, as the molecular size observed ranged between 402 Da and 3,463 Da. This is in line with the filtration cutoff of 10,000 Da used during the preparation of Imuno TF^®^, which would allow for much larger di-, tri- or polypeptides to be present in the Imuno TF^®^ extract. The peptides found in the sample were most linked to hemoglobin subunits alpha and beta. These proteins have high concentrations in the blood and can be released from red blood cells (erythrocytes), which are lysed during the purification and isolation process.

The most prominent protein acting on the immune system based on concentration in the sample and function is Talin-1. Talin-1 has been one of the first proteins shown to be recruited to the immunological synapse formed between T lymphocytes and APCs to facilitate T cell activation.(Klann et al., 2017) Supported by this function, transfer factors may mediate indirectly the innate immune response.

The addition of this set of peptides to the Reactome Knowledgebase(Fabregat et al., 2017) allowed for the identification of possible 17 pathways to which TF components could play a role. Specifically for the immune system, the identified theoretical pathways were observed for both innate and adaptative responses, as well as for the cytokine signaling: the endoplasmic reticulum (ER)-phagosome, the receptor-type tyrosine-protein kinase (FLT3) signaling, the interleukin (IL)-17 signaling, the non-canonical NF-κB pathway and the mitogen-activated protein kinase (MAPK) pathway.

The impact of TF on the phagocytic activities of the immune system was already reported,(Krishnaveni, 2013) then the ER originated phagosomes can explain, at least partially, how antigens from intracellular pathogens such as virus can be presented by MHC class I molecules.(Desjardins, 2003) The role on NF-κB was also reported as an inhibitory effect.(Salazar-Ramiro et al., 2018) If those effects are confirmed, a role in immunoregulation could be accounted to TF – for example, through the effect on IL-17, which mediates inflammation for microbial clearance, but if overstimulated can promote immunopathology.(Amatya et al., 2017) MAPK and FLT3 are also involved in inflammation and regulation of dendritic cells, respectively.(Kazi & Rönnstrand, 2019; Zhang & Liu, 2002)

In addition, two associated proteins were uncharacterized. The existence of these proteins was predicted *in silico* and no functional studies have been conducted thus far. Therefore, further studies to investigate their biological function, possibly on the immune system, are to be considered. Together with the characterization of Imuno TF^®^, its safety was also evaluated. Under the conditions evaluated in this study, Imuno TF^®^ did not present mutagenic potential at the concentrations of 5, 1.67 and 0.56 mg mL^-1^ in the systems studied, with and without metabolization, therefore it is considered safe in terms of possible interactions with the cell’s DNA. In addition, reviewing the acute oral toxicity results, it can be concluded that Imuno TF^®^ at 2,000 mg kg^-1^ bw dose can be classified as GHS Category 5 (“not classified”, according to the Globally Harmonized System of Classification and Labeling of Chemicals, GHS),(OECD - Organisation for Economic Co-operation and Development, 2001) with an LD_50_ cut-off value in female Wistar rats of 5,000 mg kg^-1^ bw (classification goes from category 1, higher toxicity, to category 5, non-detectable toxicity). The recommended oral dose of Imuno TF^®^ ranges from 50 to 100 mg, daily (equivalent to 0.7 to 1.4 mg kg^-1^ bw for a 70 kg weighing adult). This implies that Imuno TF^®^ is safe to use at the recommended dose.

## 5. CONCLUSION

TF isolated from porcine spleen (Imuno TF^®^) can be characterized as a complex set of oligo- and polypeptides and is a low fat, saturated fatty acids-free and low sugar food supplement which contains vitamins and minerals. It can be considered safe for oral use and presents no mutagenic potential. Finally, its composition accounts for possible multiple mechanisms on the immune system pathways.

---

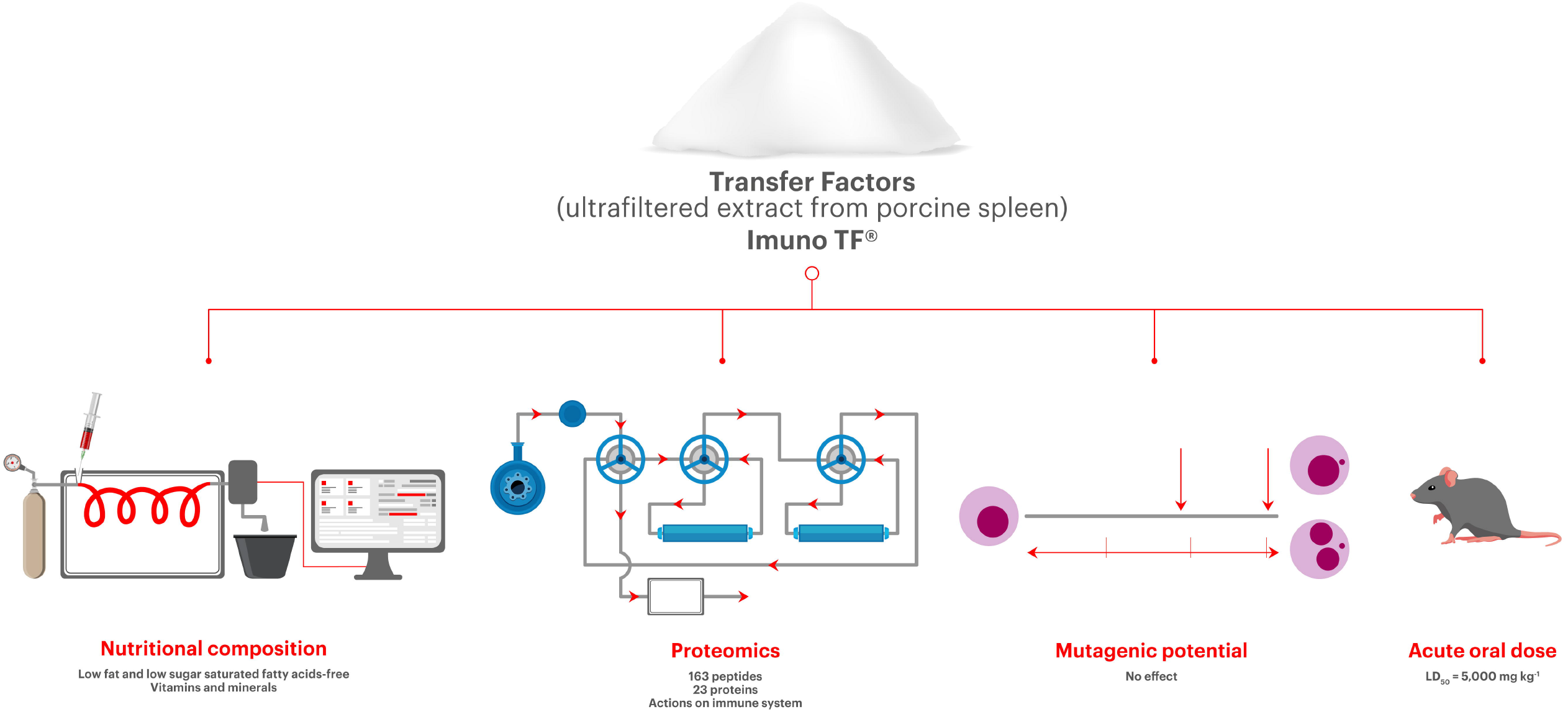

